# Bet-hedging, seasons and the evolution of behavioral diversity in Drosophila

**DOI:** 10.1101/012021

**Authors:** Jamey S. Kain, Sarah Zhang, Mason Klein, Aravi Samuel, Benjamin L. de Bivort

## Abstract

Organisms use various strategies to cope with fluctuating environmental conditions. In diversified bet-hedging, a single genotype exhibits phenotypic heterogeneity with the expectation that some individuals will survive transient selective pressures. To date, empirical evidence for bet-hedging is scarce. Here, we observe that individual Drosophila melanogaster flies exhibit striking variation in light-and temperature-preference behaviors. With a modeling approach that combines real world weather and climate data to simulate temperature preference-dependent survival and reproduction, we find that a bet-hedging strategy may underlie the observed inter-individual behavioral diversity. Specifically, bet-hedging outcompetes strategies in which individual thermal preferences are heritable. Animals employing bet-hedging refrain from adapting to the coolness of spring with increased warm-seeking that inevitably becomes counterproductive in the hot summer. This strategy is particularly valuable when mean seasonal temperatures are typical, or when there is considerable fluctuation in temperature within the season. The model predicts, and we experimentally verify, that the behaviors of individual flies are not heritable. Finally, we model the effects of historical weather data, climate change, and geographic seasonal variation on the optimal strategies underlying behavioral variation between individuals, characterizing the regimes in which bet-hedging is advantageous.

## INTRODUCTION

Understanding how organisms succeed in the face of fluctuating environmental conditions is a major challenge. One intuitive solution is phenotypic plasticity – an organism adjusts its phenotype in direct response to the current environmental condition, such as leaf size in response to lighting conditions (Sultan, 2000). However there are limitations to this approach, such as metabolic cost and the speed with which an organism can change its phenotype. Organisms can also survive changing conditions by having diversified phenotypes as a result of genetic variation; this also allows organisms to readily evolve/ adapt to new conditions. However, if the environmental changes are transient, then adaptation via heritable mechanisms will always lag behind the selective pressures. A third possible solution to the problem of uncertainty is to utilize a bet-hedging strategy (also called risk-spreading), in which developmental stochasticity produces a distribution of adult phenotypes. In diversified bet-hedging, a single genotype displays a range of phenotypes, guaranteeing that a fraction of the population is well suited to any environmental condition (Hopper, 1999; Simons, 2011; Levy et al., 2012). This strategy comes at the expense of the population also containing some poorly-adapted individuals. An elegant example is the timing of seed germination (Cohen, 1966). If all the seeds from a desert plant germinated after the first rain of the season, they would be vulnerable to extinction if there is an extensive drought before the second rain. Conversely, if the seeds all germinate later in the season, they will be at a disadvantage relative to other seeds that had germinated at the first opportunity (in typical seasons without an early drought). Thus, an optimal strategy may be for the plant to hedge its bets and have a fraction of seeds delay germination while the others respond to the first rain.

Fruit flies are one of the most studied organisms for many aspects of biology, including the basis of behavioral diversity. We chose to study this problem using the fly’s positional response to thermal gradients (thermotaxis) and asymmetric illumination (phototaxis), which are robust behaviors. The thermotactic and phototactic responses of *Drosophila* depend on a wide range of environmental and stimulus parameters (Dillon et al., 2009), such as humidity (Waddington et al., 1954), directionality of the light source (Rockwell and Seiger, 1973), and agitation state of the flies (Lewontin, 1959; Rockwell et al., 1975; Seiger et al., 1983). The type of phototactic response is particularly sensitive to the state of agitation. In most *Drosophila* species, agitated animals exhibit “fast phototaxis” toward the light source, while unagitated animals exhibit “slow phototaxis” toward shade. The former response is thought to reflect a predator evasion instinct to move skyward (Scott, 1943), while the latter reflects a thermoregulatory and anti-desiccation instinct during rest (Pittendrigh, 1958).

We found surprising variation in the slow phototactic and thermotactic responses of very recently domesticated *Drosophila* from Cambridge, Massachusetts, USA. Some individual flies strongly preferred to rest in shade (or warm regions), others strongly preferred light (or cool regions). We wondered whether this behavioral diversity represented a bet-hedging strategy to maximize fitness in the face of fluctuating seasonal or weather conditions. In order to compare the performance of bet-hedging versus a strategy in which the individual behavioral preferences are heritable (i.e. adaptive tracking *sensu* Simons, 2011) we developed a model incorporating our behavioral data with local weather and climate data from historical records. (Phenotypic plasticity in response to environmental fluctuations is unlikely to explain the behavioral differences we observed between individuals reared in essentially identical lab environments; under phenotypic plasticity, we would expect animals to adopt similar behaviors as their response to a similar environment, but this is not what we observe. Our scope here is to specifically consider a head-to-head comparison of bet-hedging and adaptive tracking strategies, both of which remain plausible explanations of the observed behavioral variation.)

We find that the bet-hedging strategy generally outcompetes adaptive tracking. Since the generation time of *Drosophila* is short relative to the seasons, seasonal temperature fluctuations can induce genetic adaptations in the spring which then decrease fitness in the summer. This reversal of selective pressures throughout the year renders adaptive tracking counterproductive. The alternative bet-hedging strategy is particularly valuable when there is high fluctuation in temperature throughout the season. Adaptive tracking is preferred, however, during seasons that are consistently warm or cold throughout, because it allows adaptation by natural selection. Interestingly, since global climate change will bring about an increase in mean temperatures, we predict that the optimal strategy will change in approximately 100 years, and adaptive tracking will become more advantageous than bet-hedging.

## RESULTS

### *Drosophila* exhibit more behavioral variability than expected by chance alone

Thermal experience has dramatic effects on the life history of *Drosophila* (Ashburner et al., 2005; Ashburner, 1978; Miquel et al., 1976). Individuals can control this experience through a variety of behaviors (Parry, 1951; Digby, 1955) including shade-seeking phototaxis and direct positional response to thermal gradients. Thus, the net resting behavior of flies will greatly affect the amount of heat they experience across their lifetime, and consequently their vulnerability to unusual weather, season and climate fluctuations. The light versus shade and thermal gradient resting preferences of animals can be readily quantified in laboratory experiments. We sought to directly measure the slow phototactic and thermotactic response of recently domesticated *D. melanogaster* flies, and assess to what extent there was individual-to-individual variability in this behavior. An isofemale line (“CamA”) was established from a single fertilized female caught in Cambridge, Massachusetts, USA.

To assess phototaxis, age-and sex-matched CamA adults, cultured on standard fly media, were assayed individually in our “slow photobox” (Figure 1A), where their light versus shade preference was measured by automated image analysis 24 times per fly (Figure 1B), once every 10 minutes. We tested 219 individuals in total, and found that their average light-choice probability was 0.32, indicating a preference for resting in the shade. The observed distribution of light-choice probabilities was considerably overdispersed compared to what would be expected if all animals were choosing the light with identical probabilities of 0.32 (*p*=4×10^−6^, 1×10^−11^ and <0.001 by Kolmogorov-Smirnov (KS) test, *χ*^2^ test of variance and bootstrap resampling respectively; Figure 1C), indicating considerable individual-to-individual behavioral variability. These results are similar to our previous findings on agitated phototaxis where we observed significant individual-to-individual variability that was not explainable by differences in age, sex, reproductive status, birth order, social interactions, or previous exposure to light (Kain et al., 2012).

**Figure 1.**
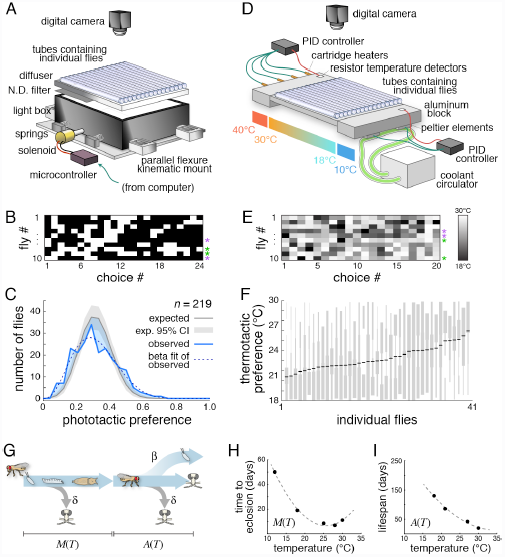
Measurement of phototactic and thermotactic variation and a model of their effect on fitness. (A) Schematic of the “slow photobox” – a device for the high-throughput characterization of slow phototaxis. Animals were placed individually into clear tubes with a lit and shady side. Their position in the tube was recorded by a camera. (B) Example of data from the slow photobox. Each row represents an individual fly’s phototactic preferences at 24 instances, spaced at 10 minute intervals. White boxes indicate lit choice and black boxes indicate shaded choice. Purple and green asterisks indicate examples of shade-and light-preferring individuals, respectively. (C) Observed histogram of the phototactic preference across individual flies (blue line). Dashed gray line indicates a best-fit beta-binomial distribution for the observed data. Gray line indicates expected distribution for the same flies if they were each to choose light with identical probabilities. Gray shaded region indicates 95% confidence interval of the expected distribution given sampling error. Shaded blue areas indicate discrepancies between the observed and expected histograms consistent with behavioral heterogeneity. (D) Schematic of the “slow thermobox.” (E) Example data from the slow thermobox, as in B. Grayscale indicates thermotactic preference over time. Purple and green asterisks indicate examples of cool-and warm-preferring individuals, respectively. (F) Histograms of thermotactic preference values across all trials (vertical, grey) for individual flies, sorted by mean preference (black bars). Black bars indicate individual mean preferences. (G) Diagram of the fly life history model, see description in text. β: birth rate, δ: death rate, *M*: metamorphosis time, *A*: adult lifespan, *T*: temperature. (H) Time to eclosion plotted as a function of temperature, as used by the model. Data points from (Ashburner 1978). (I) Lifespan plotted as a function of temperature, as used by the model. Data points from (Miquel et al., 1976).

To assess thermotaxis, similarly cultured animals were tested individually on a linear thermal gradient (Ryu and Samuel, 2002) ranging from 30°C to 18°C (Figure 1D), which spans most of the range of flies’ natural environment. The position of each of 41 flies within this gradient was measured 20 times per animal, once every 10 minutes, with their position indicating their per-trial thermal preference (Figure 1E). We observed considerable inter-individual variation in mean thermal preferences (*p*<10^−6^ by 1-way ANOVA on fly identities; Figure 1F).

### A model to compare adaptive tracking and bet-hedging strategies

Could the observed behavioral individuality represent a species-level bet-hedging strategy of assuring that the population always contains some animals well adapted to the current weather fluctuation? To test this, we proposed a model of fly development and reproduction (Figure 1G) in which an individual animal’s behavior could be treated either as perfectly inherited from the mother (i.e. adaptive tracking – AT), or as non-heritable/stochastic variation indicative of a bet-hedging strategy (BH). The model does not consider the comparison between populations with and without variability, but takes phenotypic variation as a given, based on our experimental results. Holding the magnitude of variation constant, we can evaluate which is more advantageous, adaptive tracking or bet-hedging, and under what conditions.

In considering how thermal preference might affect fitness, we recognized that the metamorphosis time from egg to adulthood depends on the temperature experienced during that period, in a relationship determined by previous experimental work (Ashburner et al., 2005; Ashburner, 1978]), with flies developing fastest at 25°C (Figure 1H). The expected total lifespan of flies also depends on temperature (Miquel et al., 1976), with flies living considerably longer at cooler temperatures (Figure 1I). We assume that the effective temperature experienced throughout adulthood depends on the integrated results of many behavioral choices for each individual fly. By contrast, we assume the temperature experienced during growth from egg through pupa depends on the thermal preference of each fly’s mother (the alternative, that developmental temperature depends on progeny preference, yields qualitatively identical results). These are clearly simplifying assumptions – the total amount of thermal energy integrated across a lifespan and the choice of oviposition site depend on more behaviors than just phototaxis and thermotaxis. But, constraining the model with empirical data on these behaviors allows us to investigate their roles in fitness. We lastly assume that throughout metamorphosis and adulthood flies face a constant risk of death (by e.g. predation, disease, fly swatter, etc.), and after reaching adulthood, flies produce new offspring at a constant rate. Thus, thermal choice represents a tradeoff for the fly: warm-preferring animals will have the benefit of faster development at the cost of shorter lifespan, the kinetics of which are temperature-dependent.

In order to formulate a single variable representing the diversity of thermal experience due to all dimensions of behavioral variability, we compared our phototactic and thermotactic observations. The effect of phototactic preference on thermal experience depends on the temperature difference between shade and sunlight. This in turn depends on numerous factors, including weather conditions, latitude, season, wind, substrate composition and duration of exposure to the sun. We measured this directly and determined that a 7°C difference between sun and shade was attained quickly after exposure to sunlight on both natural and artificial substrates in Cambridge, Massachusetts, USA. This estimate is well within the range of previous estimates of the temperature difference between insects in sunlight vs shade (Parry, 1951). We observed that the mean light-choice probability of flies in the slow phototaxis assay was 0.32, with a standard deviation of 0.13 (Figure 1C), implying a standard deviation of thermal experience of 0.89°C. The mean observed thermotactic preference was 22.7°C with a standard deviation of 1.4°C. These two observations are in agreement that individual flies experience substantial differences in thermal experience. For the model, we let the thermal preference of individual flies, integrated across all behaviors determining thermal experience, vary as an index (*T*) of 0 to 1 corresponding to a 7°C temperature difference. This corresponds to the phototactic data, which is conservative compared to the direct thermotactic measurements.

The model contains two unknown parameters, the lifelong risk of death from causes other than old-age (*δ*), and the birth rate at which new eggs are laid by sexually mature flies in the wild ( ). We have no empirical data from which to assert these values, but the behavior of the model constrains them under two reasonable assumptions – 1) that the population size of flies is the same at the end of each season as the beginning, and 2) that the mean thermal preference of the population is the same at the end of the season as the beginning, i.e. they are adapted to average conditions. These assumptions constrain the random death probability of flies in the wild to 0.013-0.044/day, and the birth probability to 0.037-0.11/mother/day (depending on what weather model is used. See Experimental Procedures for details); both of these ranges seem plausible.

### Bet-hedging outperforms adaptive tracking

We simulated a stochastic (agent-based) implementation of this model (Figure S1), tracking 100 individual flies experiencing the average seasonal temperature fluctuations (NOAA Climate Normals 2013) of a typical fly breeding season in eastern Massachusetts, USA, lasting approximately from April 1 to October 31 (Figure 2). We implemented two versions of the model. 1) For the adaptive tracking strategy (AT; Figure 2A), the thermal preference of new flies equaled that of their mother. 2) For the bet-hedging strategy (BH; Figure 2B), the preference of each new fly was drawn at random from a beta distribution fitting the observed behaviors (mean thermal preference = 0.32 and standard deviation 0.13; Figure 1C). The initial population of all simulations also followed this distribution, irrespective of strategy.

**Figure 2.**
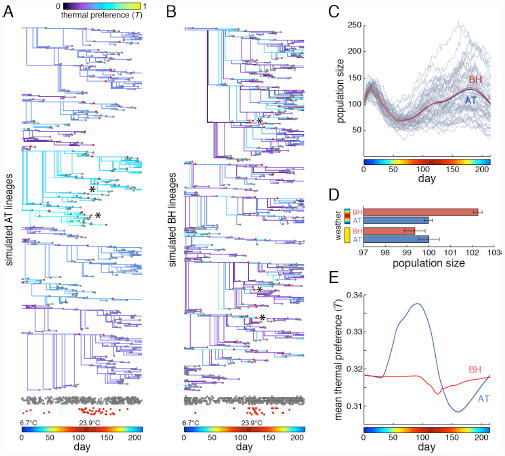
Performance of the bet-hedging and adaptive tracking versions of the stochastic model. (A) Subset of simulated lineages from one run of the model, under the AT strategy. Branch points indicate the birth of new flies; colors indicate thermal preference; gray dots indicate death events for reasons other than old age (due to parameter *δ*); red dots indicate death events due to old age. Rows of dots at bottom are projected from above for comparison, with random y-scatter for visibility. Asterisks indicate old age death events associated with high summer temperatures in light-preferring lineages. Temperature at each day is indicated by the colored bar here and in all other panels. (B) As in (A), but for the BH strategy. (C) The mean performance of a bet-hedging (BH) (red line) and adaptive tracking (AT) (blue) version of the model over time. Gray lines represent a sampling of 100 individual simulated seasons. (D) Mean final population size produced by each version of the model for either constant average weather (yellow) or seasonal weather (colored bar). Error bars are +/-1 one standard error of the mean; *n*=40,000 simulations per group. (E) Mean thermal preference of the population over time for each version of the model. Shaded regions (barely wider than plot lines) are +/-1 one standard error of the mean.

On average, the BH strategy outperformed the AT strategy by just over 2% (Figure 2C, D, *p* < 0.0001 by *t*-test), an effect that is completely absent (and non-significantly reversed, *p* = 0.64 by *t*-test) in simulations of constant seasonal temperatures. The reason for the better performance of the BH strategy is evident in an inspection of the average thermal preference of the fly population across the breeding season (Figure 2E). (The average preference changes even under BH due to temperature-dependent shortening of the lifespan of warm-seeking individuals). In the AT strategy, the cool spring selects for warm-preferring flies because their progeny will develop to maturity more quickly. However, at the onset of summer, the selection is reversed in favor of cool-preferring flies, which have a longer overall lifespan. Once the direction of selection switches, the BH strategy begins to outperform the AT strategy, because AT responds to even transient selective pressures by shifting the population mean.

### Individual phototactic preference is not heritable

The model establishes that bet-hedging is plausible explanation for the behavioral diversity seen experimentally in thermal preference. However, if the observed individuality we see truly represents bet-hedging, then the differences in preference between individual flies are probably not due to genetic polymorphisms or trans-generational epigenetic effects, which would be heritable. This hypothesis generates two predictions: 1) reducing genetic diversity by inbreeding a polygenic stock should have no effect on the breadth of its behavioral distribution, and 2) the progeny of light- (or shade-) preferring parents should exhibit the same distribution of behaviors as the entire parental generation, not their specific parents. (These predictions were tested in the phototactic paradigm because of its higher throughput and our use of its parameter values in the model). We compared the behavioral distribution of our polygenic isofemale CamA line with that of the line “inbred-CamA” which was inbred by sibling matings for 10 generations. Inbreeding had no significant effect on the mean or variance of the behavioral distribution (Figure 3A). Using inbred-CamA we set up multiple crosses comprising a male and a virgin female that both either prefer the shade or the light (Figure 3B, C). If their individual photopreferences are due to genetic polymorphisms between flies, then their progeny should have a correspondingly shifted mean photopreference relative to the original population. However, we found there was no difference in the mean photopreferences of broods derived from shade-preferring parents versus light-preferring (Figure 3B–E); thus, heritable polymorphisms determine at most a small component of each individual’s behavior, consistent with a bet-hedging strategy. Moreover, the distributions of brood photopreferences were indistinguishable from the parental distribution, in variance as well as mean (Figure 3D,E).

**Figure 3.**
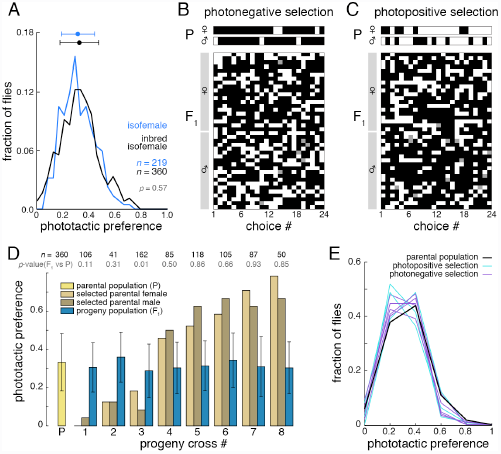
Individual phototactic preference is not heritable. (A) Observed histogram of the phototactic preference across individual CamA flies (blue) and inbred-CamA flies (black). Points and bars represent the distribution mean and +/-1 standard deviation. *p*-value from KS test. (B) Representative samples of the phototactic scores of a shade-preferring male and female (top) and the phototactic scores of their resulting progeny (bottom). Each row represents an individual fly’s phototactic preference over time. White boxes indicate lit choice, black boxes indicate shaded choice, and gray boxes a missing value. (C) As in (B), but for light-preferring parents and their progeny. (D) Phototactic indices for strongly biased shade-or light-preferring individuals (tan and brown bars) and their resultant progeny (dark blue bars). The dashed line and yellow bar indicate the original pool of animals from which strongly biased individual parents were selected. Numbers above bars indicate sample size, with *p-*values from KS test uncorrected for multiple comparisons. Error bars are +/-1 one standard deviation. (E) Histograms of phototactic preferences within the respective progeny (D).

### An analytic version of the model yields similar results

The heritability intrinsic to the AT strategy means that in a finite population simulation (such as in our model population of 100 virtual flies; Figure 2) the mean thermal preference of the population can vary significantly from replicate to replicate due to the stochastic nature of the model (Figure 2C). AT may lock in maladaptive thermal preferences due to drift, and the rate at which this happens depends critically on the simulated population size (Wright, 1931). Since it was arbitrary to simulate 100 animals, and effective population sizes in the wild are unknown (and perhaps far too large to simulate efficiently (Karasov et al., 2010)), we developed an analytic version of the model, in which the population size was effectively infinite and immune to stochastic effects. In this implementation, sub-populations of flies with specific thermal preferences were determined by a set of difference equations (see Experimental Procedures). The analytic versions of the BH and AT strategies performed similarly to the simulations of individuals (Figure 4A, B), with BH outperforming AT by 1.1% by the end of the summer, and the AT model undergoing two selective sweeps of opposite direction.

**Figure 4.**
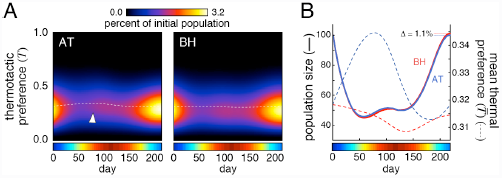
Performance of BH and AT using an analytic implementation of the model. (A) Abundance of flies as a function of thermal preference and time for AT and BH strategies under the analytic model. Arrowhead indicates adaptive thermal positivity during the spring. Dashed white line indicates the mean thermotactic preference.(B) Population size (solid lines) and mean thermal preference (dashed lines) over time of BH (red) and AT (blue) versions of the analytic model.

### Incorporating historical weather data into the model

To test the effects of daily temperature fluctuations and cloud cover, we ran the analytic model against historic weather data collected in Boston, Massachusetts, USA (NOAA Climate Normals 2013) (Figure 5A). The temperature in each day of the simulation was taken from actual historical data from that day, on a year-by-year basis. Cloud cover was implemented by assuming that the temperature difference available for flies to respond to (i.e. between sun and shade) each day was proportional to the average cloud cover of that day. Not surprisingly, reducing the temperature difference available to flies (due to cloud cover) reduced the magnitude of the advantage of the BH strategy (to around 0.2% for years 2007-2010) (Figure 5B). We initially thought that short-term heat waves (or cold spells) might be enough to confer an advantage to bet-hedging, but these were found to make little difference. In 2010 the BH advantage was lowest. The weather that year was consistently warmer than in the others, particularly in the spring and fall (Figure 5C), exerting a comparatively uniform selective pressure for cool-seeking, thereby reducing the advantage of bet-hedging.

**Figure 5.**
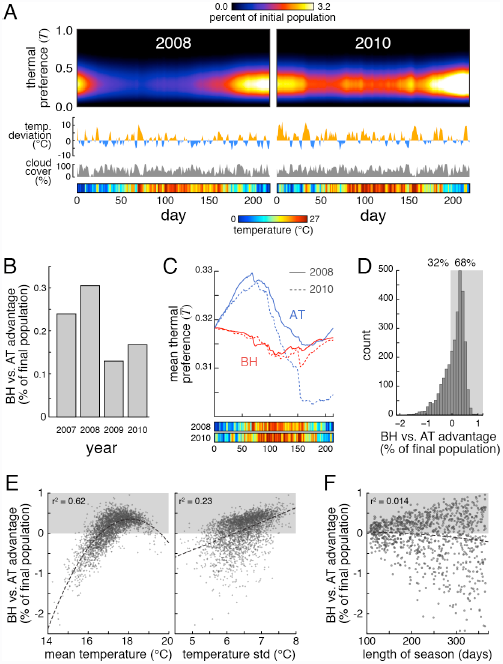
BV vs AT in historical and modeled breeding seasons. (A) Abundance of flies as a function of thermal preference and time in the BH version of the analytic model, applied to historical weather data (temperatures and cloud cover) from 2008 and 2010. Orange and blue traces indicate temperature deviation from daily normals. Gray traces indicate daily cloud cover percentage. Colored bars indicate the daily mean temperature. (B) BH versus AT advantage as a percent of the final population vs. year ((popBH-popAT)/popAT*100). (C) Mean thermal preferences for AT (blue) and BH (red) versions of the real weather analytic model for 2008 (solid lines) and 2010 (dashed lines). Colored bar as in (A). (D) Histogram of BH versus AT advantage as a percent of final population using the analytic model across 3000 simulated seasons. Shaded region indicates the simulations in which BH outperformed AT. (E) Scatterplot of BH versus AT advantage versus mean temperature (left panel) or the standard deviation of the temperature (right panel), across 3,000 simulated seasons. Shaded region indicates the simulations in which BH outperformed AT. *r*^2^ values reflect quadratic fits (dashed lines). (F) Scatterplot of BH versus AT advantage versus breeding season length, across 1000 simulated weather seasons. Shaded region and *r*^2^ value as in E.

### Mean temperature and temperature range are most predictive of the BH vs AT advantage

We developed statistical models of the daily temperature fluctuations and cloud cover that allowed us to simulate realistic random breeding seasons, and systemically tested the factors favoring the BH and AT strategies. Across 3000 random seasons, BH outperformed AT 68% of the time (Figure 5D). We examined numerous metrics describing the simulated seasons (Figure S2) and found two in particular that were predictive of the magnitude of the BH vs AT advantage (Figure 5E): the temperature mean and standard deviation. BH outperforms AT when the season has a typical temperature while exceptionally hot or cold seasons favored the AT strategy. Additionally, AT performs poorly during “intense” seasons – those with cold springs and falls, and hot summers, because of strong opposing selective sweeps.

We also analyzed the effects of shorter or longer breeding seasons by compressing or stretching random temperature and cloud cover histories into seasons ranging from 107 to 365 days (Figure 5F). The average relative advantage of BH versus AT did not depend on season length, however the variance of BH advantage increased with season length. Only long seasons exhibited strong advantages for either BH or AT, presumably because increasing the number of generations per season increases the potential for adaptation, whether it be productive or counter-productive.

### Global climate change is predicted to shift evolutionary strategy from BH to AT

Across the 3000 random seasons, the BH vs. AT advantage never exceeds ~1% per season, but could drop as low as ~-2% in some seasons (Figure 5D). Despite the longer negative tail in this distribution, the small advantage of BH over AT in most summers quickly accumulated across simulations of multiple sequential seasons (Figure 6A), indicating this strategy was highly favored on longer timescales. However, we found that an increase of only 2°C to the mean seasonal temperature was sufficient to change the evolutionary dynamic in favor of adaptive tracking (Figure 6B). Conservative models of global climate change predict i (Meehl et al., 2007). Thus, while seasonal weather fluctuations generally favor bet-hedging in thermal preference behavior, climate change will likely cause a phase-shift in the evolutionarily optimal strategy toward adaptive tracking.

**Figure 6.**
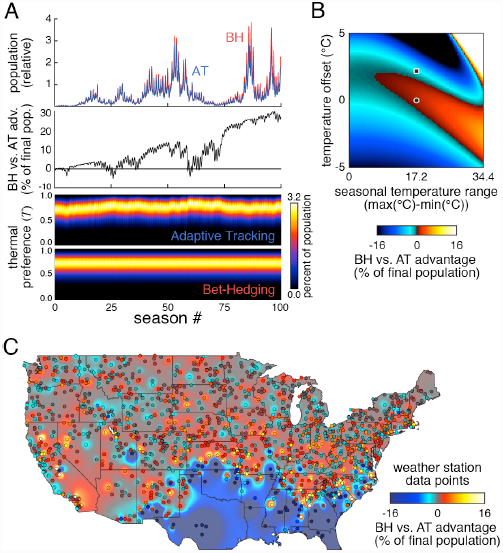
Climatic and geographic variation in BH vs AT advantage. (A) Relative population sizes for the BH (red) and AT (blue) versions of the model (top) and cumulative BH vs. AT advantage (middle) over the same 100 random simulated seasons. Bottom panel shows the corresponding abundance of flies as a function of thermal preference and time, across 100 seasons, for each strategy. (B) Phase space of BH vs. AT advantage as a function of the two most predictive metrics. Color indicates magnitude of the advantage. Circle indicates current state while the square indicates the state if the average temperature were to increase 2°C. (C) Geographic map of BH vs AT advantage. Data points correspond to specific NOAA weather stations; background coloration is interpolated. See Supporting Information for details.

### Geographical variation in BH vs AT advantage

Lastly, we considered to what extent the BH vs AT advantage we saw with Boston weather data was location specific. We ran the model using mean daily temperature data from more than 1400 weather stations across the continental United States (NOAA Climate Normals 2013) and compared the performance of the BH and AT strategies (Figure 6C). Our model predicts substantial regional variation in the optimal strategy. In most locations, BH maintains a small advantage. In the deep south, where the breeding season is year-long, allowing more time for adaptation, BH performance is much worse than AT. However the temperature extremes and shortened breeding season of regions just north (or at higher elevation in the southern Appalachians) renders BH strongly advantageous. This is consistent with the observation that long breeding season can strongly favor either AT or BH (Figure 5F). Consistently, the short breeding seasons of higher latitudes and the Rocky Mountains favor neither AT nor BH strongly. AT appears to be favored along the Pacific coast, which is characterized by low temperature fluctuations.

## DISCUSSION

Here we explored whether a bet-hedging strategy could explain the large observed variation in thermal preference in *Drosophila*, as measured in phototactic and thermotactic paradigmsseasonal temperature selective pressures, adaptive tracking is always lagging; by the time the population has adapted to the cool springtime with increased warm-preference, summer arrives. Therefore, it is better that the behavioral preference of individual flies be non-heritable so that there are always spring-adapted and summer-adapted animals being born. The bet-hedging advantage is strongest under two conditions. 1) Highly variable temperatures (cool springs coupled to hot summers) magnify the selective pressure on the adaptive tracking group and thus produces larger counterproductive sweeps as the temperature fluctuates throughout the season. This is consistent with the observations of seasonally fluctuating allele frequencies in flies (Bergland et al., 2013). 2) When mean temperatures are atypical, the BH strategy is disadvantaged by not being able to evolve. In one example, the year 2010 was warmer on average and its spring was particularly warm, reducing the seasonal temperature variability. Both of these factors gave the AT strategy a relative boost for being able to evolve and thus reducing the overall BH advantage (Figure 5B,C).

Beyond adaptive tracking and bet-hedging, another major strategy for dealing with environmental heterogeneity is plasticity, in which organisms adaptively tune their phenotype in direct response to environmental fluctuations. The set of plasticity strategies can even include hybrid strategies such as the moment-to-moment regulation of the extent of bet-hedging in response to environmental conditions. In the absence of constraints, such as metabolic cost or limits on achievable phenotypes, a plasticity strategy is tautologically optimal (DeWitt and Langerhans, 2004), though such constraints surely exist. Generating an empirical estimate of the costs imposed on *Drosophila* in response to environmental fluctuations. We observed striking behavioral variation in populations of animals grown in essentially identical conditions (laboratory culture); to first approximation, there were no environmental fluctuations (e.g. variations in ambient temperature or luminance) to which a plasticity strategy could respond. Second, under conditions of convex fitness functions (i.e. with a single predominant mode of fit phenotypes), plasticity strategies can be at a disadvantage compared to bet-hedging strategies even if they come with low costs (DeWitt and Langerhans, 2004). The unimodal relationships between temperature and eclosion time and lifespan (Figure 1H) yield a convex fitness function in our case, suggesting that plasticity may be outcompeted by bet-hedging (or even adaptive tracking), even if it comes at a relatively low cost.

Our analysis focused on *Drosophila melanogaster*, a species with a relatively short reproductive cycle capable of producing several generations within the breeding season. It is likely that species generating fewer generations per season (i.e. K-selected species) would be less subject to the pitfalls of an adaptive tracking strategy since they would respond less to any temperature fluctuation. While our model did not permit us to realistically change the life history of our simulated *Drosophila* in the context of real weather data, we were able to simulate changes in the length of the breeding season (Figure 5F). Shorter seasons are comparable to a K-selected life histories because they yield fewer generations per season. We found that, as hypothesized, shorter seasons reduce the difference between adaptive-tracking and bet-hedging strategies, while long seasons can favor either strategy depending on other factors (i.e. Fig 5E).

This modeling highlights the importance of population-level properties, namely the amount of variation and the heritability of that variation. Population-level traits touch on the topic of group selection (Wilson and Wilson, 2008), and indeed aspects of bet-hedging were sometimes conflated with group selection in the literature (Hopper, 1999). However, our models do not directly address this controversial issue because they have no reliance on specific population structures, (the concept of which largely evaporates when considering non-heritable traits). Importantly, selection still operates, in all implementations of our model, at the level of the individual.

Two avenues for future investigation emerge from our results. First, flies captured and assayed at different time points throughout the season should show differences in their mean thermal preference (Figure 2E), that reflect their mode of inheritance. Specifically, flies using a AT strategy and caught in the early summer would be comparatively warm-seeking, while flies using a BH strategy would be comparatively cool-preferring at the height of the summer, when the high temperature selectively shortens the lifespan of warm-seeking individuals. However, analysis of behavior across the breeding season must consider seasonal changes in allelic frequencies (Bergland et al., 2013). Second, flies from locales with large seasonal weather changes (e.g., Boston, Massachusetts, USA) may have greater behavioral variation than those from milder, less variant climates (e.g., coastal central California, USA; Figure 6C).

There is also a third prediction from these models concerning the affect of climate change on these strategies. Due to incrementally increasing mean temperatures over time, adaptive tracking becomes a better option as the organisms continually adapt to the new normal. An increase of 2°C will be sufficient to favor adaptive tracking over bet-hedging, a change predicted to take approximately one hundred years. While it is speculation to predict how a population would respond to such a switch, we note that both phototactic (Dobzhansky and Spassky, 1969) and thermal preference (Dillon et al., 2009) are heritable in outbred populations, presumably under genetic control. Heritability in these behaviors is a prerequisite for evolving the heritability of BH vs AT strategies themselves. It is plausible that a switch in selective pressure on strategies could increase adaptive tracking by favoring lineages with deeper developmental canalization.

The conclusions drawn from the models here are not meant to say that bet-hedging is the sole explanation for behavioral variation. However, we have found that under the constraint of experimental data on the magnitude of behavioral variability between individuals, and with a minimal set of assumptions, bet-hedging appears to be a more adaptive explanation of behavioral variation than deterministic genetic heterogeneity. Indeed, we believe that real *Drosophila* probably utilize at least three strategies – bet-hedging, adaptive tracking, and phenotypic plasticity – to optimize its survival in an uncertain world.

## EXPERIMENTAL PROCEDURES

### Behavior

The *Drosophila melanogaster* line CamA was established from a single mated female caught from the wild in Cambridge, MA USA and propagated in the lab for approximately two generations at typical *Drosophila* culture densities prior to behavioral testing. The line inbred-CamA was derived by 10 generations of sibling matings. All flies were cultured on standard growth medium (Scientiis) in 25°C incubators at 30-40% relative humidity on a 12-12h light-dark cycle. We found no difference in the behavioral responses of males versus females and merged their data.

Age-and sex-controlled flies were placed singly into tubes in the “slow photobox,” which is illuminated from below by diffused white LEDs (5500K, LuminousFilm) (Figure 1A). A 50% neutral density filter was used to generate a lit half and shaded half for each tube. The rig is mounted on kinematic flexure mounts allowing ~1cm translation parallel to the testing tubes, under the control of a solenoid/microcontroller system driving vibration at 20 Hz. Agitation of the animals induced them to run and thereby reset their position between successive measurements of their light/shade preference. Each trial consisted of agitation (three 2s pulses, each separated by a 1s pause), an interval of 577s, acquisition of the photo used to score animal position, and a 15s interval completing the 10m trial. Animal position was determined by subtracting the background image of the rig and calculating the centroid of all pixels that had changed relative to the background (on a tube-by-tube basis), subject to a noise-eliminating threshold.

The slow thermobox was fabricated by placing the acrylic tray of choice tubes used in the slow photobox down on a slab of aluminum with thermal grease. The aluminum slab was in contact with two larger aluminum blocks, one warmed to 40°C with resistive heating elements, and one cooled to 10°C with thermoelectric coolers (Peltier elements). The temperature of both larger blocks was held constant by PID controllers reading insulated resistor temperature detectors (3-wire, 100ohm). The 30-18°C gradient achieved within the choice tubes was measured using a infrared thermometer gun and was highly linear. For each of 20 trials, animals were first agitated by flowing air into the choice tubes, dislodging the animals toward the warm end. After 9.5 minutes the tubes were photographed and the position of each animal measured digitally.

### Temperature measurement

Temperature differences between sun and shade were measured using an infrared thermometer gun on partly cloudy days in the summer and autumn. In one set of of comparisons we measured the temperature of substrates in the shade of clouds, and then waited until ~5 minutes after the cloud had passed and measured their temperature in sunlight. In another set of comparisons, we compared adjacent sunlit and shaded (e.g. by a building or road sign) substrates of the same orientation. Measured substrates included grass, brick, pine branches, tree bark, gravel etc.

### Raw data and code

All raw data used in this study, as well as all code used for data acquisition, statistical analysis and modeling are available at http://lab.debivort.org/bet-hedging-seasons-and-the-evolution-of-behavioral-diversity-in-Drosophila.

### Statistics

Data from individual flies that did not move upon agitation for 3 or more successive trials were discarded since these measurements were clearly non-independent from trial to trial. Sequential slow phototactic choices were found to have an average of 0.054 bits of mutual information across individuals, indicating effective independence (0 bits indicates complete independence in every animal, 1 complete dependence). Therefore we modeled the expected distribution of light-choices with a binomial distribution with parameter *p* equal to the average light-choice probability of all animals tested, and parameter *n* equal to the number of trials, 24.

### Modeling

See Results, Figure 1D, and Figure S1 for a description of the model. In the bet-hedging implementations of the model, each fly was randomly assigned a thermal preference drawn from the experimentally observed preference distribution (fit by a beta distributions; Figure 1C). In adaptive-tracking implementations, the seed population was initialized in that way, but all subsequent animals were assigned a preference identical to their mother’s preference. Stochastic simulations of finite populations were seeded with 100 flies with ages uniformly distributed on [*M*(*T*), *A*(*T*)] – respectively the mean ages of eclosion and death due to old age – since flies may over-winter as adults (Izquierdo, 1991). We also implemented a version of the model in which the seed population was synchronized to the egg stage. This model was qualitatively indistinguishable. Flies in this initial population were assigned to have developed at random in the sun versus the shade with a probability equal to the population mean thermal preference. Individual flies were simulated, removed from the virtual population at random according to the parameter *δ*, and born stochastically at a rate *β* from mature flies already in the population. To determine the temperature experience by each fly each day, its thermal-choice preference was multiplied by the temperature difference between light and shade and added to the shade temperature. The birth and death rate parameters were identified (by grid search or hill-climbing algorithm) as the unique pair of values that satisfy two assumptions: 1) the fly population neither grows nor diminishes across the breeding season, i.e it is at numerical equilibrium, and 2) the mean thermal preference does not evolve across the breeding season, i.e. flies are adapted to typical conditions. For every distinctive weather model, parameter fitting was independently performed using the adaptive tracking implementation. See Table S1 for parameter values. The analytic version of the model was implemented analogously using a system of difference equations, but could efficiently be used to evaluate historical and simulated daily temperature deviations and cloud-cover values (see Supporting Information).

## ACKNOWLEDGEMENTS

We thank Chris Stokes for help with fabricating the slow photobox and we thank Julien Ayroles, Sarah Kocher, Greg Lang and Sean Buchanan for helping analyze the data and model. This work was supported by the Rowland Junior Fellows Program.

## Supplementary materials

### Modeling

In the analytic implementation of the model, clouds reduced the maximum ambient temperature difference attainable by individual flies in proportion to the mean daily cloud cover fraction. Historical daily temperature deviations were normally distributed, and modeled using a 30 parameter autoregression filter of normally distributed white-noise. Random cloud cover was generated by drawing a season-long sequence of values from the observed (non-Gaussian) distribution of cloud cover fractions. These values were then shuffled until the new cloud cover sequence was no longer correlated with the original sequence (*r*<0.1), under the constraint that the autocorrelation of the simulated sequence was correlated to that of historical cloud data with *r*>0.998, thus preserving temporal statistical structure of the sequence. Historical cloud and temperature deviation data were uncorrelated (*r*=0.02), so simulated sequences of these variables were derived independently.

The analytic model was implemented using the following system of difference equations:

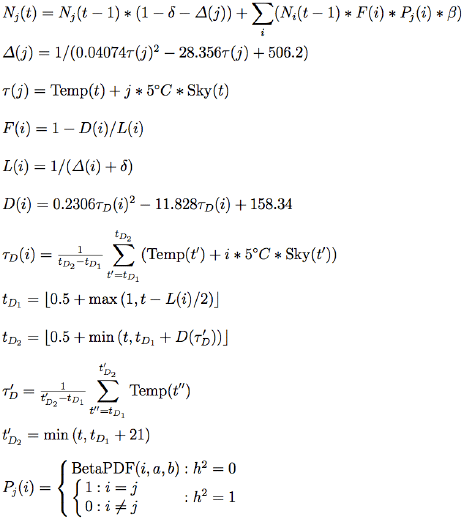

Here, *N*_*j*_(*t*) is the number of flies alive at time *t* with thermal preference *j*. *Δ*(*j*) is the rate at which flies die due to old age as a function of *j*. τ(*j*) is the effective temperature experienced by flies with thermal preference *j*, with Temp(*t*) indicating temperature and Sky(*t*) indicating respectively the temperature and cloud cover fraction at time *t.* The summation term in *N*_*j*_(*t*) indicates the number of flies born at time *t* with thermal preference *j*, born from parents with thermal preference *i*, which depends on the population sizes of flies with thermal preference *i* at time *t*−1(*N*_*i*_(*t−l*)), the fraction of each of those parental subpopulations which are fertile (*F*(*i*)) and the probability densities of parental thermal preference (*P*_*j*_(*i*)) conditioned on the thermal preference of the progeny (*j*), and given the alternative BH vs AT strategies. (*Pj*(*i*) is coded as a matrix with probability entries in row *j*, column *i.* For strategy H, it is the identity matrix; for strategy B, every row of *P*_*j*_(*i*) equals the beta-fit distribution from Figure 1C.) *F*(*i*) depends on the ratio of development time *D*(*i*) to total lifespan *L*(*i*) of flies with thermal preference *i*. *D*(*i*) depends on the effective temperature experienced by parents (as this determines egg laying site) during development τ_*D*_(*i*) which we approximate as the mean effective temperature across a range starting at time *t* minus half the typical lifespan, and ending *D*(τ’*D*) days later (bounded by the time endpoints of the simulation). Development time is dependent on integrated temperature, which in turn depends on the length of development, given temperature’s temporal fluctuation. So the calculation of *D*(τ’*D*) reflects one level of recursion in the calculation of this feedback. τ’*D* is calculated as the average temperature from t’*D1* through 21 days later, an interval approximating half a typical lifespan. The results of the analytical model are very robust to the choice of the intervals in this recursion approximation, as well as the number of recursive levels implemented.

In simulations of sequential seasons, the mean thermal preference of the initial population of each season was set to the mean of population at the end of the previous season, but the variance was reset to match the empirical data. In geographic simulations, breeding seasons were defined as all days between the first day of the year in which temperatures reach 6.5°C and the first day when mean temperatures fall below 10°C, the same thresholds used in the Boston season. The non-parity in these values reflect our understanding that the first thaw suffices to end diapause while the first frost is sufficient to trigger it. The specific predictions associated with some stations are sensitive to these bounds, but the overall geographic patterns are not. The *β* and *δ* parameters were fit independently for each station automatically using a hill-climbing algorithm. Included stations were chosen at random from the 7500 stations in the NOAA data set, however, the algorithm was unable to fit the model parameters for some stations in very hot regions, i.e. some of the deep south and the Mojave desert, so station geographic sampling is not unbiased. Background interpolation in Figure 6C was done pixel by pixel using the function 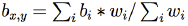, i.e. the average of all stations indexed by *i* and weighted by *w*_*i*_, where 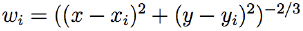, i.e. inverse Euclidean distance from the pixel (*x*,*y*) to station *i* raised to the third power. This exponent was chosen to ensure a sharp drop-off with distance from the stations, but is otherwise arbitrary.

**Figure S1.**
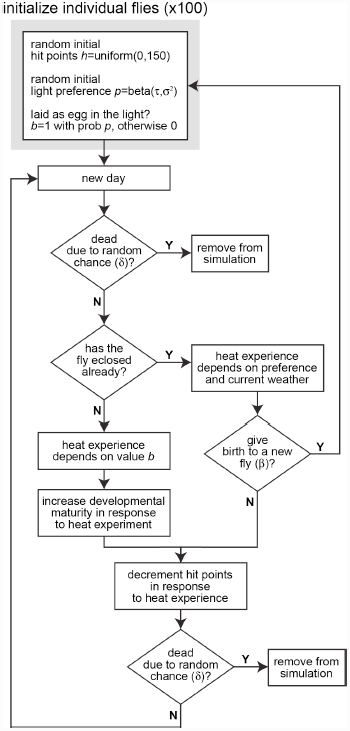
Flowchart of the stochastic agent-based implementation of the fly life history model. See Results for additional explanation

**Figure S2.**
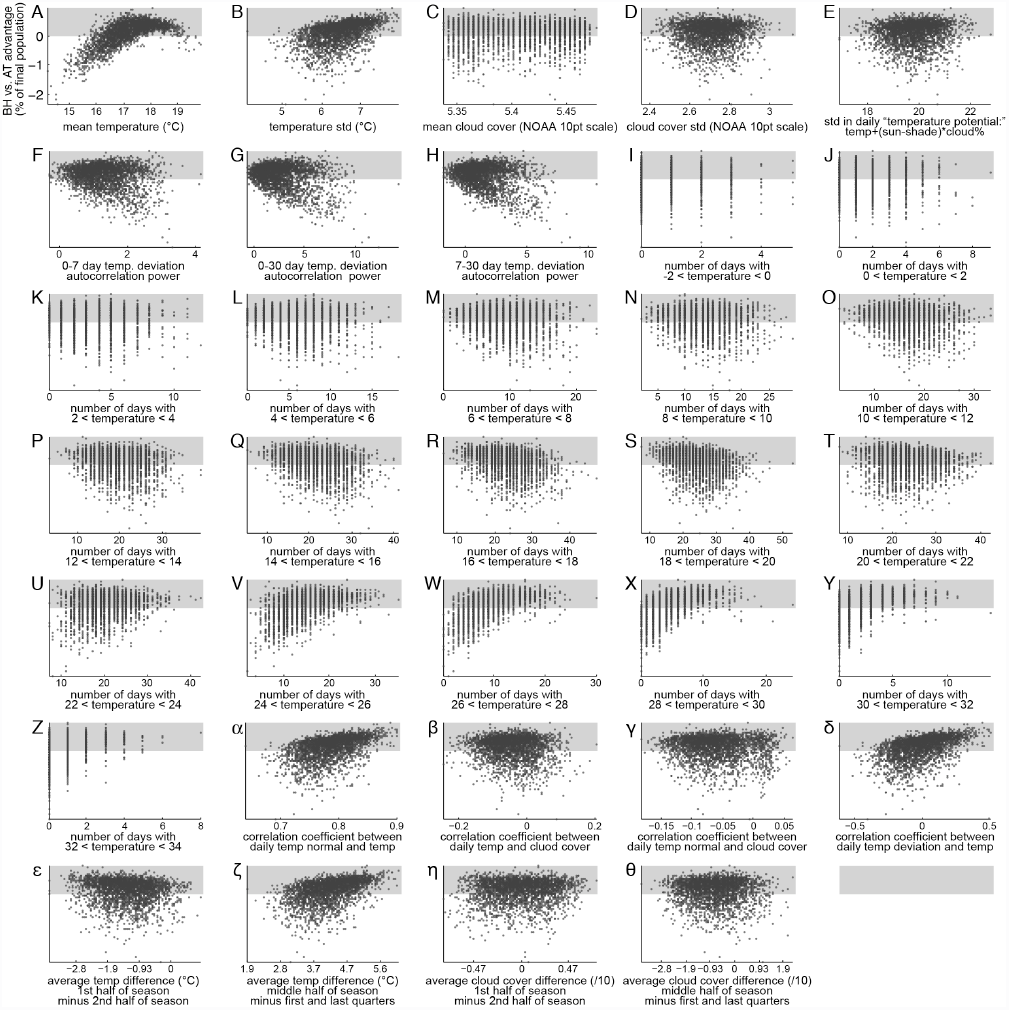
BH versus AT advantage versus various seasonal measures. All Y-axes as in first panel. F-H were calculated by summing the normalized autocorrelation vector of the daily temperature deviations for the specified range of offsets. Most measures showing clear relationships with BH-AT advantage reduce to either mean temperature (A) or temperature standard deviation (B). As examples: Seasons with many days of moderate temperature (R, S) correspond to seasons of low temperature standard deviation. Conversely, seasons with more very high temperature days (X-Z) correspond to hot years (A). Seasons with greater correlation between daily temperature and temperature normal (α, γ) have more extreme temperature ranges, corresponding to (B).

**Table S1.**
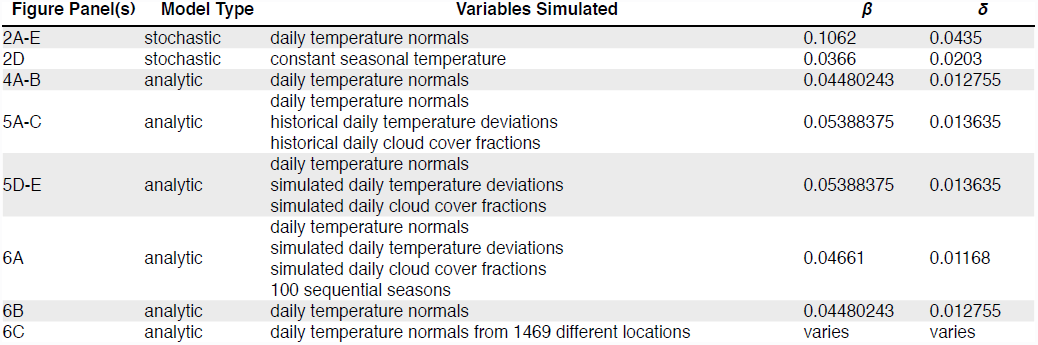
Variables simulated in each implementation of the model and values of the fit birth and death rate parameters.

## Abbreviations

AT: adaptive-tracking
BH: bet-hedging
ANOVA: analysis of variance
NOAA: National Oceanographic and Atmospheric Administration
LED: light-emitting diode
PID: proportional-integral-derivative

